# Automated Brain Masking of Fetal Functional MRI

**DOI:** 10.1101/525386

**Authors:** Saige Rutherford, Pascal Sturmfels, Mike Angstadt, Jasmine Hect, Jenna Wiens, Marion van den Heuval, Dustin Scheinost, Moriah Thomason, Chandra Sripada

**Affiliations:** Department of Psychiatry, University of Michigan, Ann Arbor, MI; Department of Electrical Engineering and Computer Science, University of Michigan, Ann Arbor, MI; Merrill Palmer Skillman Institute, Wayne State University, Detroit, MI; Department of Cognitive Neuropsychology, University of Tilburg, Tilburg, The Netherlands; Department of Radiology and Biomedical Imaging, Yale University School of Medicine, New Haven, CT; Department of Child and Adolescent Psychiatry, New York University Langone, New York, NY

**Keywords:** fetal, fMRI, functional imaging, brain segmentation, deep learning, convolutional neural network, automated processing, open-source software

## Abstract

Fetal resting-state functional magnetic resonance imaging (rs-fMRI) has emerged as a critical new approach for characterizing brain development before birth. Despite rapid and widespread growth of this approach, at present we lack neuroimaging processing pipelines suited to address the unique challenges inherent in this data type. Here, we solve the most challenging processing step, rapid and accurate isolation of the fetal brain from surrounding tissue across thousands of non-stationary 3D brain volumes. Leveraging our library of 1,241 manually traced fetal fMRI images from 207 fetuses (gestational age 24-39 weeks, M=30.9, SD=4.2), we trained a Convolutional Neural Network (CNN) that achieved excellent performance across two held-out test sets from separate scanners and populations. Furthermore, we unite the auto-masking model with additional fMRI preprocessing steps from existing software and provide insight into our adaptation of each step. This work represents an initial advancement towards a fully comprehensive, open source workflow for fetal functional MRI data preprocessing.

## 1 Introduction

Resting-state functional magnetic resonance imaging (rs-fMRI) has emerged as a powerful tool for studying development of the brain’s network architecture. In recent years, this methodology has been applied to the human brain in utero, opening a window into a period of functional development that was otherwise inaccessible. Studying fetal fMRI has the potential to illuminate the nature and manner in which the brain’s network architecture is initially assembled, affording powerful new insights into neurodevelopmental origins [1, 2, 3, 4]. Despite this potential, progress has been slow due, in part, to the lack of image analysis tools tailored for fetal imaging data. Though many tools and software packages exist for fMRI analysis, these tools were designed with adult and child data in mind and encounter specific problems when applied to fetal functional data. In particular, these tools are not set up to handle encasement of the head inside a heterogeneous tissue compartment, high degree of variation in image properties across subjects, and the extensive position adjustments made by the fetus during a typical scanning session.

Progress made towards improving fetal MRI methodology can broadly be divided into two fields: image acquisition and image post-processing. Image acquisition improvements have occurred mainly with respect to fetal structural MRI, particularly in anatomical (primarily T2 HASTE; Half-Fourier Acquisition Single-shot Turbo spin Echo imaging) and diffusion tensor imaging (DTI) [5, 6]. Advances have been made in inter-slice motion correction and volume reconstruction [7, 8], mapping structural connectivity [9, 10, 11, 12, 13], comparing different MRI signals [14]. These strategies have made it possible to use sparse acquisition sequences, which alleviate movement concerns [15, 16], and enable more sophisticated analytic approaches, such as morphometric [7, 17, 18, 19], cortical folding [20], and cytoarchitectural examinations [21]. In contrast, papers focusing on image post-processing improvements are markedly few and have also primarily focused on structural imaging. Papers suggesting possible solutions for the analysis of fetal functional MRI data are rare [14, 22]. In this work, we focus on methods for improving image post-processing for fetal functional MRI data.

Challenges associated with analysis of fetal fMRI have been discussed in a growing number of studies [23, 24, 25, 4] and reviews [26, 27, 28]. These works have focused on image characteristics: motion, size of the fetal brain, susceptibility artifacts introduced by surrounding maternal tissues, and physiological noise of both mother and fetus. Previous work has highlighted important areas for development, but to our knowledge, no one has proposed a preprocessing pipeline for fetal fMRI and released in the open science framework.

The most time-consuming step in preprocessing fetal fMRI is differentiation of the fetal brain from the surrounding maternal compartment at each acquisition time point. This is achieved by generation of an exemplar mask that marks all in-brain voxels. This mask is critical for the entire preprocessing pipeline and for subsequent activation and/or connectivity analyses. Tools developed to segment the adult brain, such as the Brain Extraction Tool (BET) from FSL [29] and 3dSkullstrip from AFNI [30] are not effective in generating exemplar masks in fetal imaging because the surrounding tissue is more complex and the fetal brain is not in a standard orientation, making it more challenging to identify. As a result, previous studies have relied on manual generation of brain masks [31, 23, 24, 25, 4]. While manual methods are tedious and time consuming, to date, they have been necessary to achieve acceptable standards.

Here, we present an automated approach to the problem of fetal brain segmentation from surrounding tissue. Leveraging a large corpus of manually traced human fetal fMRI masks, we trained a convolutional neural network (CNN) to replace this labor-intensive preprocessing step. CNN’s are a powerful tool for effectively identifying complex, non-linear patterns in spatially structured high-dimensional datasets [32]. They are increasingly utilized in image processing applications in both medical and non-medical settings [33, 34]. In the context of fetal brain segmentation, prior work has investigated the application of CNN’s to segment fetal structural T2-weighted volumes [35, 36, 37, 38, 16, 39, 40, 41]. These models, however, were developed to segment the fetal brain from anatomical images, and do not translate to functional time series data. Compared to structural data, functional data is significantly lower resolution and, due to movement, requires a larger quantity of individual segmentations. Here, we extend prior work by developing and validating a tool for automatically segmenting the fetal brain from functional MRI data. Ultimately, we connect our auto-masking model with an automated version of a previously manual preprocessing workflow.

All code discussed in this paper, along with a protocol, is available on GitHub (https://github.com/saigerutherford/fetal-code). This pipeline embodies an initial set of guidelines for fetal fMRI preprocessing, and associated protocols are expected to evolve over time through improvements by the user community in response to new knowledge and innovations in the field.

## 2 Project Background

Primary data used for pipeline development were acquired at Wayne State University School of Medicine during the course of projects supported by National Institutes of Health (NIH) awards MH110793 and ES026022. These projects aimed to characterize the development of functional neural systems beginning in utero in relation to prenatal exposures and future neurobehavioral development. Participants were recruited from obstetric clinics located within the Detroit Medical Center. Eligible participants were at least 18 years of age, assessed as having uncomplicated, singleton pregnancies, and had no contraindications for MRI. A physician (with a clinical relationship to eligible patients) initiated contact, and those expressing interest in participating were then introduced to a member of the research team who explained study procedures and answered patient questions. All participants provided written informed consent before undergoing MRI examination. The study protocol was approved by the Human Investigation Committee of Wayne State University.

## 3 Methods

### 3.1 Participants and Data

Resting-state functional MRI was obtained from two cohorts, Wayne State University (WSU) and Yale University. WSU cohort consists of 197 fetuses (gestational age 24-39 weeks, M=30.9, SD=4.2). Twenty-one of these fetuses were scanned at two time points in utero. Both time points are included in this study, however, they are counted as a single subject. WSU fetal MR examinations were performed on a Siemens Verio 3T scanner using an abdominal 4-Channel Flex Coil. Scanning protocols have evolved since the inception of the project in 2012. The majority of data were acquired using echo-planar sequence (TR/TE: 2000/30; 4mm slice thickness, axial, interleaved ascending slice order, 360 volumes) (See Supplementary Materials for all scan parameters). Multi-echo resting-state sequences were also collected in a portion of these subjects (TR/TEs: 2000/18,34,50). The Yale University cohort contains 10 fetuses scanned twice longitudinally (gestational ages 30-36 weeks, M=32.7, SD=1.9). The Yale scanner was a Siemens Skyra 3T using a 32-channel abdominal coil (TR/TE:2000/30; 3mm slices, 32 slices parallel to the bi-commissural plane, 150 volumes). Due to lack of tools for automated segmentation of the fetal brain, research personnel were trained to manually draw fetal brain masks using BrainSuite software [42]. In line with prior work, in the present analysis manually generated brain masks are used to judge the accuracy of automated segmentation methods.

### 3.2 Auto-Masking

#### 3.2.1 Experimental Pipeline

Due to multiple manually drawn masks per subject, WSU data were randomly separated at the subject level into training, validation, and test sets with 129, 20, and 48 subjects (855, 102, and 206 volumes) respectively. The training set was used to optimize the model. The validation set was used to gauge the generalization performance of the network during training and to determine when to stop training. The test set is held out and used only after training was completed to evaluate the performance of the model.

For those interested in using this model on unlabeled data from potentially unseen MRI scanners, we wanted to further demonstrate the transfer ability of our model. We re-trained the CNN a second time by combining both the WSU training and test data (177 subjects/1,066 volumes), and used the same validation set (20 subjects/102 volumes) for determining when to stop training. We tested this CNN model on an additional fetal functional dataset (referred to as the Yale test set) collected in a separate population (New Haven, CT) and on a different MRI scanner. The Yale test set was comprised of 57 volumes from 10 unique subjects.

#### 3.2.2 Data Preprocessing

Minimal preprocessing was performed and included removing the image orientation from the image header, resampling and zero padding the images to consistent voxel sizes (3.5 × 3.5 × 3.5 mm) and dimensions (96 × 96 × 37).

#### 3.2.3 Network Architecture

The U-Net style CNN network architecture [43] implemented in this pipeline was motivated by prior work in Salehi et al. [41]. The architecture features repeated blocks of 3×3 convolutions followed by the ReLu activation function assembled into a contracting path, followed by an expanding path. In the contracting path every second convolution is followed by a 2×2 max pooling operation. In the expanding path, every second convolution is followed by a 2×2 upsampling operation using nearest-neighbor interpolation. Every other feature map in the contracting path is concatenated along the depth dimension to the corresponding map in the expanding path, which helps the network learn the appropriate location of the output mask. The final layer is convolved with two 1×1 filters to produce an output mask with channels equal to the number of output classes.

The network separates 3D image volumes into 2D axial slices and operates on each slice independently. We chose to implement a 2D rather than 3D network in order to reduce computational costs. This model includes steps for converting raw NIFTI images into a format readable by the network, and steps for converting the output of the network into a NIFTI-formatted, 3D brain mask.

The model was implemented using Tensorflow (version 1.4.1). Training and testing of the network were performed using a GPU, but CPU testing times are were also evaluated to provide an additional point of reference.

#### 3.2.4 Training Procedures

During training, the weights in a CNN are minimized with respect to a loss function that determines how well the network is learning from the training data. We optimized our network with respect to per-pixel cross entropy, with weights determined using the Adam Optimizer [44]. Adam is a first-order gradient method that updates the weights adaptively based on previous and current gradients. Even using an adaptive optimizer, we found that using a learning rate decay improved performance. The initial learning rate was set to 0.0001 with exponential decay rate of 0.9, applied every 10,000 batches. The model was trained until performance no longer improved on the validation set. In addition, we augmented the 2D axial slices in the training data through 90-degree rotations and horizontal and vertical flips. While augmentations were done on 2D slices, the same rotations and flips were applied to all slices in order to preserve the 3D shape. These augmentations capture the non-standard orientation of the brain in fetal volumes.

#### 3.2.5 Evaluation

The evaluation process was performed over multiple steps. First, we evaluated our network’s ability to mask the fetal brain using the dice coefficient. The dice coefficient is the most common evaluation metric for testing segmentation. It measures the percent overlap between two regions: the predicted brain region and the true brain region. It is defined between 0 and 1, where 0 means there is no overlap between the two regions, and 1 means the two regions are identical. We also report the Jaccard index, Hausdoff surface distance [45], sensitivity (true positive rate: brain voxels are correctly identified as brain), and specificity (true negative rate: nonbrain voxels are correctly identified as nonbrain) of our network on the WSU and Yale held-out test sets. Detailed mathematical definitions of these metrics can be found in the supplemental material and guidelines for the choices of these evaluation metrics are described in Taha et al [46].

After training, we calculated dice coefficients for all auto-masks that have a manually drawn complement, though we report values only for volumes in the test data, as performance within the train and validation datasets does not reflect model performance on new data.

### 3.3 Comparison to other methods

In addition, to aid in evaluation of obtained evaluation metrics, we performed a secondary analysis to demonstrate that current methods for adult brain extraction perform poorly when applied to fetal data. We used the Brain Extraction Tool (BET) implemented in FSL, 3dSkullstrip from AFNI, and the fetal anatomical U-Net from Salehi et al. (2017), enabling comparison of approach efficacy. Of note, evaluation metrics can be improved by separating testing data into challenging versus non-challenging images [41], but that approach is not favored and not used here as this diminishes the representativeness of estimates when applied across complex and varied data sets.

### 3.4 Failure Analysis

After model performance was evaluated, we conducted a failure analysis to discover patterns of intrinsic image characteristics that may influence auto-masking performance. First, we examined the relationship between the dice coefficient and gestational age. Next, we examined the effect of varying the number of augmentations in the training dataset. Finally, we evaluated whether image artifacts, brain size ratio (brain volume relative to the entire image volume), or position of the brain in the center of imaging space influence model performance by qualitatively examined images with dice coefficients below 0.9.

### 3.5 Application of the auto-masks

Auto-masks were generated for all available data, summarized in table 1. We note that the number of subjects reported in table 1 is higher than the number of subjects reported in the auto-mask model train/validation/test split. This is due to the time-consuming nature of manual brain masking and thus 51 subjects had available raw data to process but no manually drawn brain masks available. Also many subjects who did have manually drawn masks, had many more un-masked volumes. Visual inspection was used to confirm accuracy and quality of every auto-mask volume within a subject’s time-series, and a pass/fail scale was used. Auto-masks are output as spatial probability estimates, wherein voxel values equal to one correspond to highest probability of being brain. Failed auto-masks are flagged and discarded. Probability map brain masks were then clustered, thresholded, and binarized. These steps are taken in order to discard small, non-brain clusters that may have been included in the probability map brain mask. Binarized masks were then resampled back into subject native space where they were applied to the native image using an image multiplier, resulting in segmented brain volumes corresponding to each fetal fMRI data timepoint.

**Table 1:** Summary of all functional data collected at Wayne State University and preprocessed using the proposed automated preprocessing pipeline. The amount of time spent creating brain masks for all available data by automated methods is compared to the amount of time it would take to manually create brain masks.

|  |  |
| --- | --- |
| **Total number of subjects** | 248 |
| **Number of subject scanned longitudinally** | 163 |
| **Total number of visits** | 411 |
| **Total number of runs** | 777 |
| **Total number of masked volumes** | 190,056 |
| **Time to auto-mask all volumes** | 10.6 hours |
| **Time to manually mask all volumes** | 22 years |

### 3.6 Other Preprocessing

In addition to segmentation of the fetal brain from surrounding maternal tissue, fetal imaging preprocessing requires a number of additional steps: motion denoising, realignment of volumes within a time series, and group-level normalization to a fetal template. All of these steps are challenging because frame-to-frame displacement is elevated in fetal studies, and the fetal brain is typically not in a single standard orientation.

In prior studies by our group, a reference frame from each quiescent period was chosen to be masked. The mask would then be applied to every volume within the low movement period, not only the volume it was drawn on. Due to the time-consuming nature of manual masking, it was not feasible to mask every volume. A central goal of auto-masking is to be able to mask a full time-series in a fraction of the time it takes to manually draw a mask for a single volume. Realigning all volumes within a time series aids in automatically identifying low movement periods that are usable for further activation or connectivity analyses. Volumes in low movement periods are identified by the realignment parameters that are output as a text file and can also be visualized from the saved realignment plots. Typical realignment of fMRI data is done on full time series using the middle volume (in time) as a reference volume. Due to notably high movement across a fetal time series, using a single reference volume for realignment may not be an optimal approach. We selected the MCFLIRT FSL realignment tool [29]. While this tool still uses a single reference volume, it estimates a linear transformation between volume *n* and the reference volume and then uses this transform as the starting point to estimate the *n* + 1 to reference transform. These transformation matrices are applied to each volume of the full time series to produce a new data set comprised of realigned volumes. This step also produces a text file and plot that summarize the six rigid-body realignment parameters across time, which can be subsequently used in identification of motion outliers and motion censoring in later processing stages. Here, we applied the fsl_motion_outliers routine as a data-driven means of defining periods of high and low fetal movement.

After masking and realignment, time-series data are converted into a 4-dimensional file, moved into group template space (for multi-subject averaging) using linear warping, and spatially smoothed. Flexibility is built into the pipeline such that the user is able to define whether data are normalized to a common reference template, or alternatively, to age-specific fetal templates, see Serag et al [18]. A linear normalization is implemented via FLIRT [29]. After normalization, all volumes are spatially smoothed with a user-specified Gaussian kernel. Importantly, all software tools used within this preprocessing pipeline (TensorFlow, Python, AFNI, FSL) are free and open-source. All commands can be implemented in a shell script, which can be run from the command line.

### 3.7 Quality Checking

While this methodology employs fully automated techniques for preprocessing fetal resting-state fMRI data, manual quality assurance processes are necessary at key transition points throughout the pipeline. Specifically, our standard process includes initial review of raw time-series data, screened as a movie. Initial inclusion criteria are that the brain is in the field of view and unobstructed by artifacts, and that within the time series, there are periods of minimal fetal movement. We exclude data not meeting these criteria. However, the majority of Wayne State University data passes this stage because long scan durations are used, and fetuses rapidly cycle through quiescent states.

Additional steps in the quality control protocol are implemented after auto-masking, realignment, and normalization. At these stages timeseries data are again visually inspected to assure that no errors were introduced during these stages of preprocessing. Several parameters that are automatically generated during the pipeline should be used in complement to manual quality checking. These parameters include realignment parameters, motion plots, and metrics from the fsl_motion_outliers command. After the time-series has been realigned, calculating the dice coefficient between consecutive volumes could also provide a metric of data quality – a dice coefficient equal to one represents two perfectly overlapping (or realigned) images.

## 4 Results

### 4.1 Auto-masking performance

Our CNN auto-mask model learned to accurately discriminate fetal brain from surrounding structures in fetal brain fMRI images. We evaluated the model on two held-out test sets. Applied to the Wayne State University (WSU – 206 volumes from 48 unique subjects) and Yale (57 volumes from 10 unique subjects) test cohorts, the models achieved a per-volume average dice coefficient of 0.94 and 0.89, respectively. The CNN’s performance in terms of dice coefficient, jaccard coefficient, Hausdorff surface distance, sensitivity, and specificity across both test sets is summarized in Table 2. Figures 1 and 2 provide examples of agreement between manual and auto-masks in both test sets.

**Table 2:** Performance of auto-mask model and existing masking software evaluated in two independent test sets from Wayne State University (WSU) and Yale University. These values represent the mean and (sd).

|  | WSU |  |  | Yale |  |  |
| --- | --- | --- | --- | --- | --- | --- |
|  | Auto-mask | BET | 3dSS | Auto-mask | BET | 3dSS |
| **Dice** | 0.94 (0.067) | 0.22 (0.13) | 0.24 (0.10) | 0.89 (0.13) | 0.22 (0.06) | 0.25 (0.08) |
| **Jaccard** | 0.89 (0.069) | 0.13 (0.086) | 0.14 (0.07) | 0.82 (0.13) | 0.13 (0.03) | 0.15 (0.05) |
| **HD(mm)** | 12.11 (22.4) | 112.6 (36.7) | 103.3 (26.0) | 19.25 (14.5) | 95.2 (30.1) | 92.8 (24.2) |
| **Sensitivity** | 0.90 (0.04) | 0.13 (0.08) | 0.14 (0.07) | 0.84 (0.12) | 0.12 (0.03) | 0.15 (0.05) |
| **Specificity** | 0.99 (0.0007) | 0.99 (0.003) | 0.99 (0.002) | 0.99 (0.002) | 0.99 (0.004) | 0.99 (0.002) |

**Figure 1:**
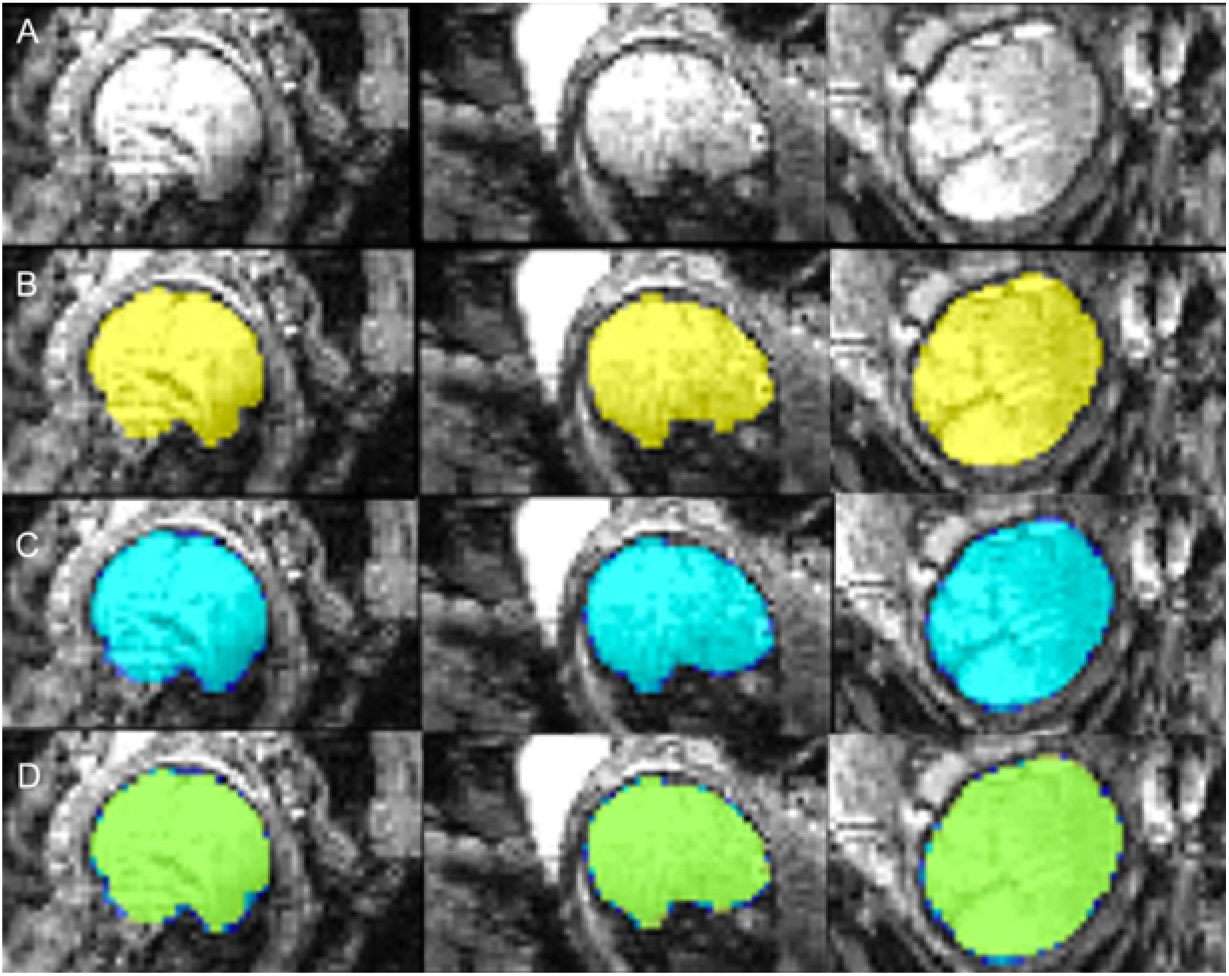
Comparison of manual and automated masks in WSU cohort. A) Raw volume; B) Hand drawn mask; C) Auto mask; D) Conjunction of hand drawn (yellow) and auto (blue) masks, overlap between hand and auto masks shown in green. Data collected in Detroit, MI at Wayne State University.

**Figure 2:**
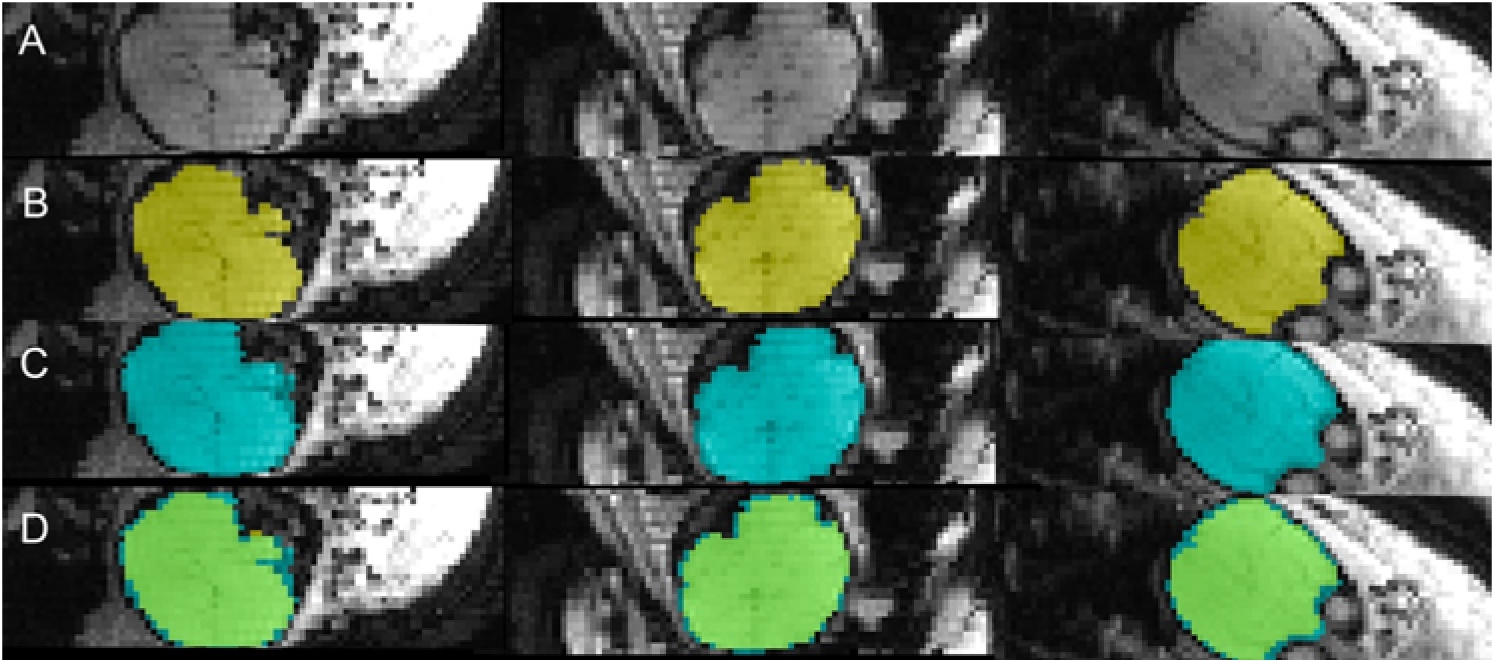
Comparison of manual and automated masks in Yale cohort. These results demonstrate the auto-mask model’s ability to transfer to data collected in different populations and/or scanners than the original data used to train the model. A) Raw volume; B) Hand drawn mask; C) Auto mask; D) Conjunction of hand drawn (yellow) and auto (blue) masks, overlap between hand and auto masks shown in green. Data collected in New Haven, CT at Yale University.

### 4.2 Comparison to adult auto-masking tools

As a point of reference for the dice coefficient achieved by our method, we performed an additional analysis using several existing auto-masking tools: Brain Extraction Tool (BET), 3dSkullstrip, and the fetal anatomical U-Net. Applied to the same test set data, as expected, these tools performed significantly worse. Evaluation metrics are reported in Table 2 and examples of the masks generated using these tools are shown in figure 3. The fetal anatomical U-Net masks were empty in most cases, and therefore the evaluation metrics failed and are not reported in Table 2. These results highlight that areas of the maternal compartment have high contrast boundaries and varied image intensity creating serious challenges for standard masking routines and severely compromise performance.

**Figure 3:**
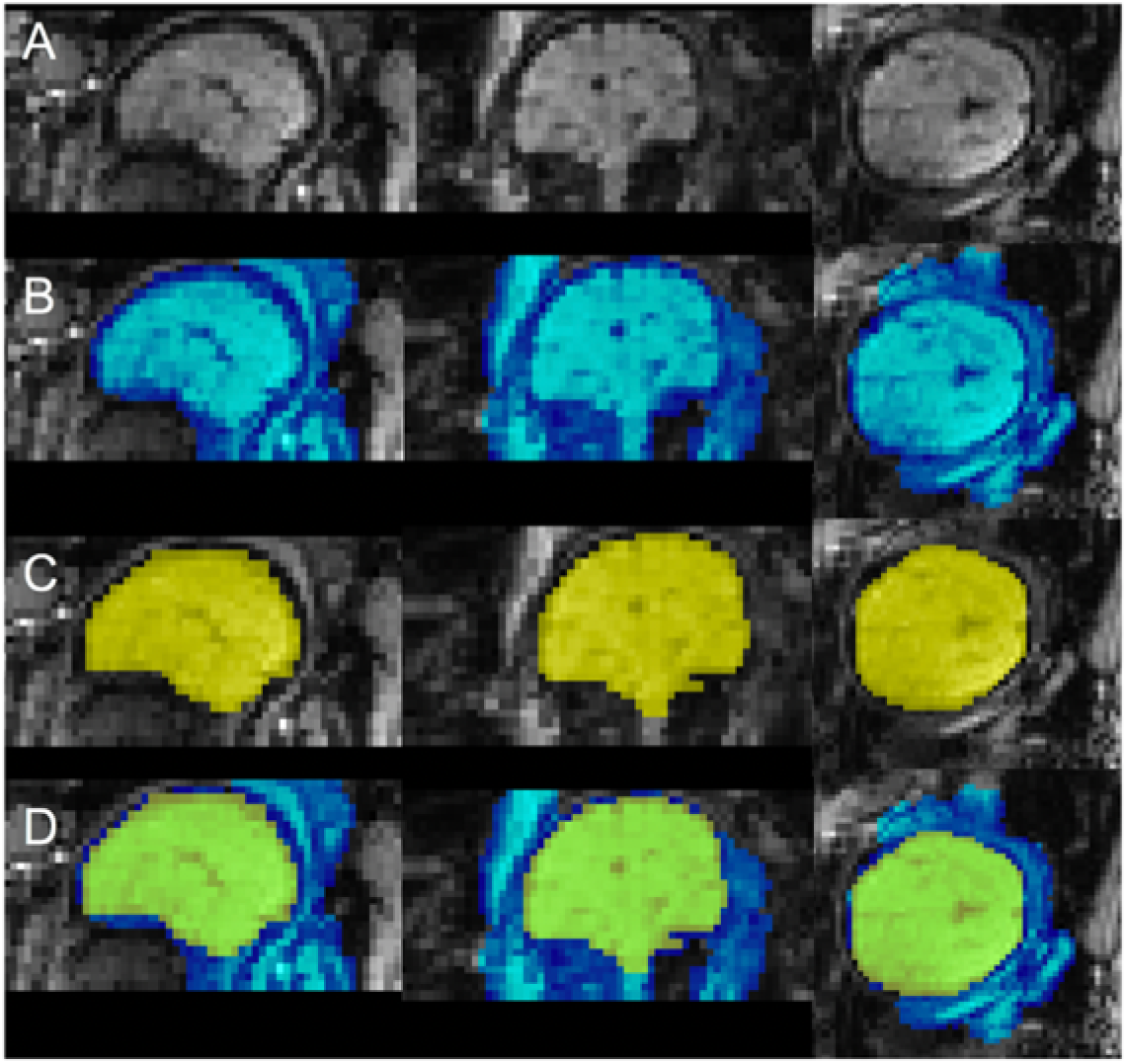
Limitations of the existing toolkit for automated brain masking. A) Raw volume; B) BET mask; C) Hand drawn mask; D) Conjunction of hand drawn (yellow) and BET (blue) masks. The BET masks do not properly capture the fetal brain’s boundary.

### 4.3 Age and Data Quality Failure Analysis

Examination of the effect of fetal age on performance of the auto-mask algorithm revealed a significant positive association between dice coefficient and gestational age (*r* = 0.14, *p* = 0.02), jaccard index and gestational age (*r* = 0.15, *p* = 0.01), and sensitivity and gestational age (*r* = 0.24, *p* =1 × 10^−3^). There were not significant correlations between gestational age and specificity or gestational age and Hausdorff surface distance. This effect is demonstrated in Figure 4. This relationship may result from older fetuses having larger brain volumes that intrinsically have higher effective image resolution. The found highly statistically significant relationships between mask volume and dice coefficient (*r* = 0.23, *p* = 2 × 10^−3^), mask volume and jaccard coefficient (*r* = 0.27, *p* = 1.53 × 10^−5^), mask volume and sensitivity (*r* = 0.26, *p* = 1.88 × 10^−5^), and unsurprisingly between mask volume and gestational age (*r* = 0.84, *p* = 1.44 × 10^−68^). We also found that significant aliasing, particularly phase wrap-around negatively impacted auto-mask performance (example shown in Figure 5), and that the algorithm also performed more poorly for images in which the brain had a large displacement from the image origin. This observation also may explain why the performance improved with an increasing number of augmentations in the training dataset.

**Figure 4:**
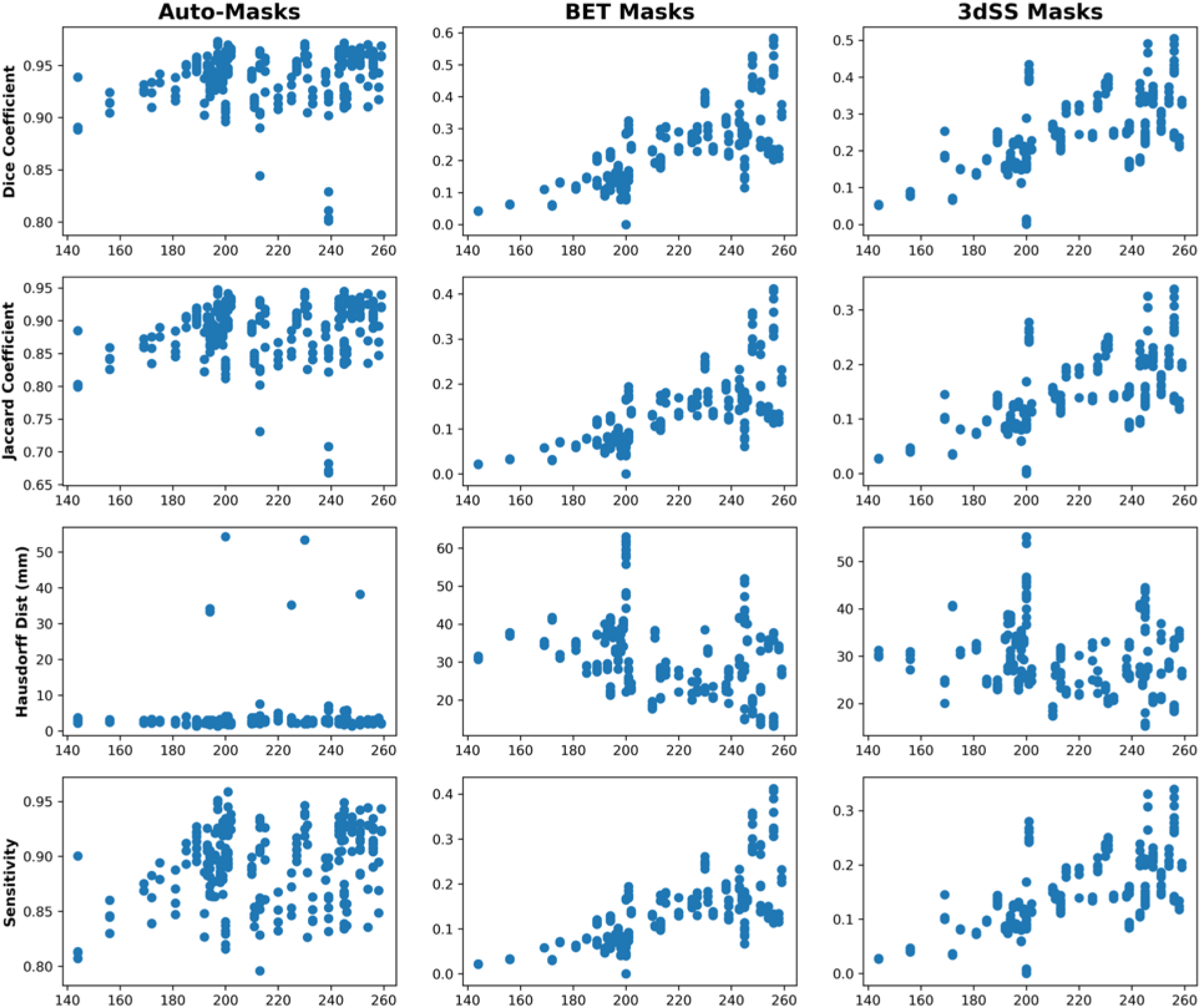
Evaluation of auto-masking model. The relationships between fetal gestational age (in days) at scan is shown on the x-axes and performance of auto-masking in the WSU test set (48 subjects, 206 volumes) y-axes. We calculated the evaluation metrics on a per-volume basis. However, the values shown here are on a per-subject basis in order to examine the relationships with age. There were significant correlations between gestational age and dice coefficient (*r* = 0.14, *p* = 0.02), jaccard coefficient (*r* = 0.15, *p* = 0.01), and sensitivity (*r* = 0.24, *p* =1 × 10^−3^). There were not significant correlations between gestational age and specificity or Hausdorff surface distance.

**Figure 5:**
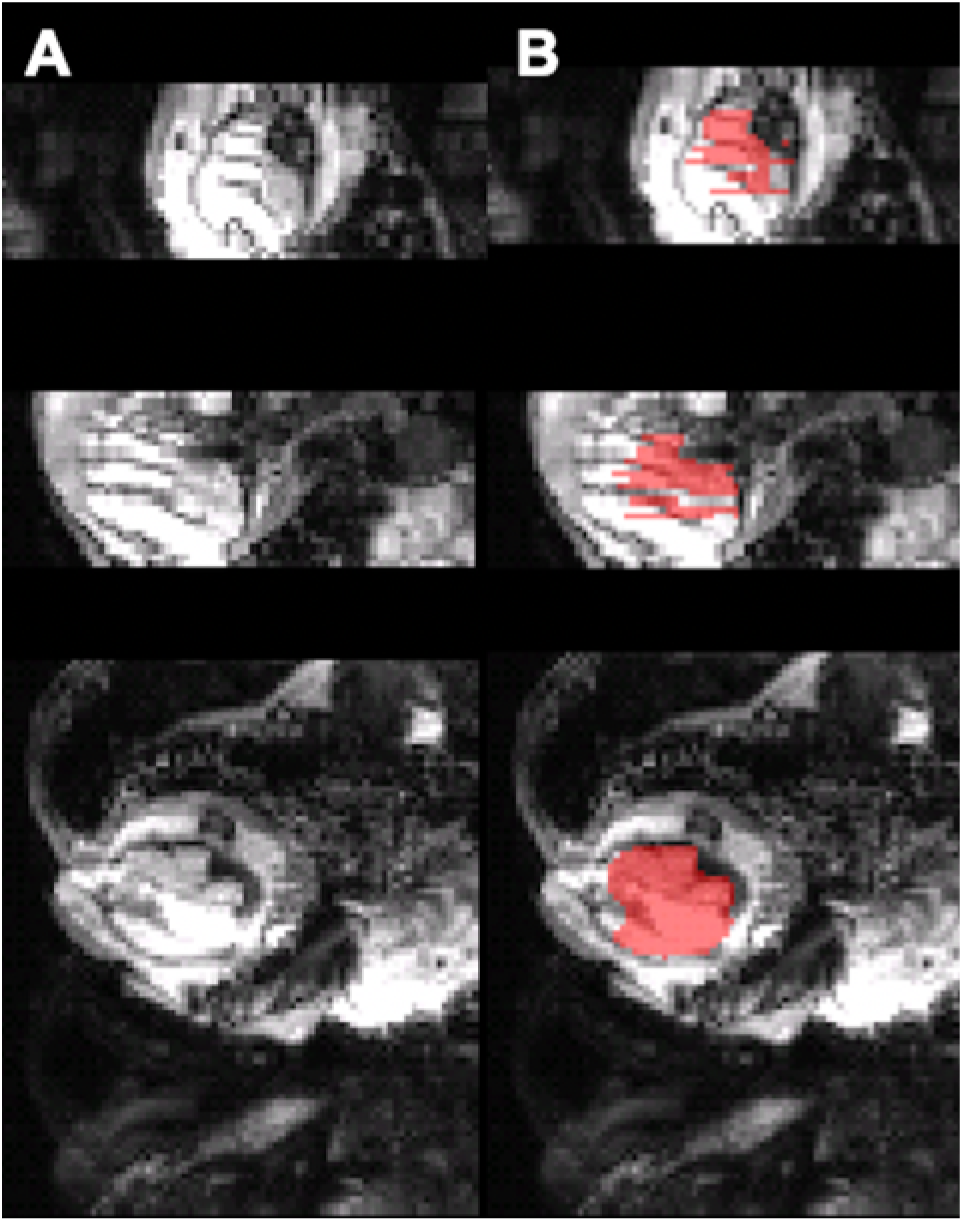
Example of auto-masking failure. A) Raw volume with severe artifacts present. B) Auto-mask that failed to capture a large portion of the fetal brain.

### 4.4 Computational Time and Hardware

An often-noted property of deep learning models is their ability to substantially surpass human speed in completing complex tasks. Our auto-masking model illustrates this acceleration. The training time refers to the wall-clock time it took our CNN model to converge to a set of weights that minimize the dice coefficient on the validation set. Training was stopped after signs of overfitting were observed, that is, performance on the validation set was no longer increasing. The total training time of the model was 3 hours and 46 minutes on a GeForce GTX 1080 Ti GPU. Testing time refers to the time it takes to run the CreateMask.py script in order to load the input raw volume (nifti file type to numpy array conversion) and output a predicted auto-mask. Additionally, we report the time to create auto-masks for all volumes within a subject’s time series. As some potential users of these preprocessing methods may not have access to GPU computing resources, we report testing time in GPU as well as CPU environments. These testing times are summarized in Table 3.

**Table 3:** Estimated computing times of all preprocessing steps. The time to run the full pipeline is for a single subject with 360 volumes. Testing GPU was a NVIDIA Quadro P6000.

|  | GPU | CPU |
| --- | --- | --- |
| **Auto-mask a single volume** | 0.2s | 2.5s |
| **Auto-mask an entire time series** | 1.2m | 15m |
| **Realignment** | - | 5m |
| **Normalization** | - | 7m |
| **Time to run full pipeline** | 13.2m | 27m |

### 4.5 Other Preprocessing

An additional benefit of our approach for auto-masking all individual volumes is that multiple realignment strategies are now possible. After the fetal brain has been extracted, the time series data can enter more typical preprocessing steps for which child/adult tools have been developed. The main difference when applying these tools is that the fetal brain is commonly in a non-standard orientation, and fetal data exhibits substantially increased head motion.

To address the first challenge, we incorporated realignment using MCFLIRT [29] into the pipeline. This method uses the preceding volume to provide an initial realignment estimate for the current volume. The idea behind this choice is that there is likely to be high spatial displacement between volumes 10 and 100 because they were collected 180 seconds apart, but less displacement between volumes 10 and 11 collected just 2 seconds apart. With regard to the problem of elevated head motion, errors introduced by movement cannot, at present, be fully corrected in fMRI time series. This fact necessitates application of stringent criteria for retaining only low-motion volumes to ensure data integrity, which is the approach taken by most studies to date [2, 3, 1, 4, 47]. We recommend two criteria for retaining low motion volumes: 1) framewise displacement less than 0.5mm, and 2) number of consecutive volumes reaching those criteria must be 10 or more. The latter rule reduces the number of breaks in the time series introduced by the data elimination scheme. These criteria await systematic testing against alternative schemes. An overview of the entire suggested preprocessing stream is provided in Figure 6.

**Figure 6:**
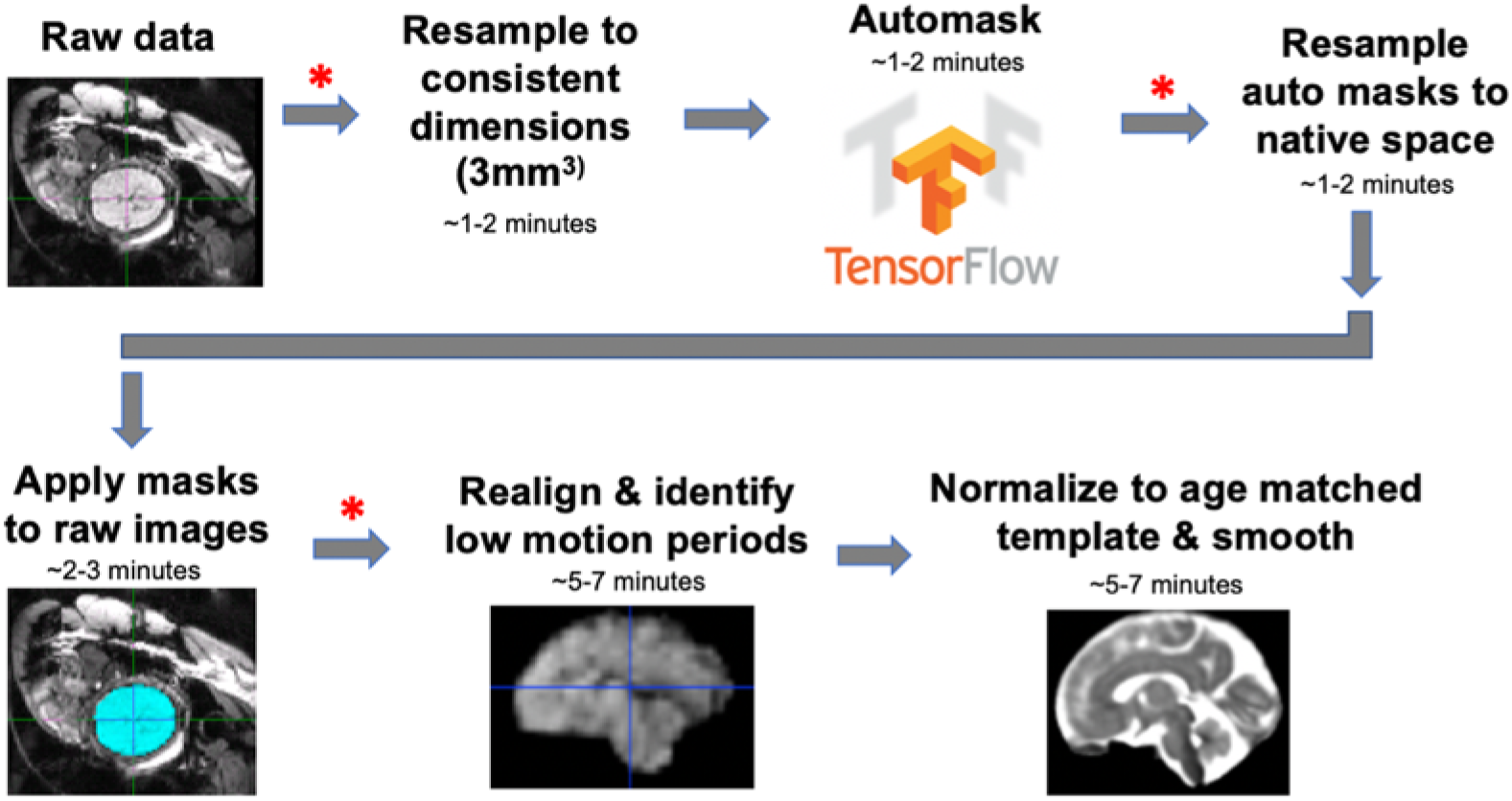
Overview of proposed processing pipeline. All steps in the proposed preprocessing stream are shown, with red asterisk representing where visually quality checking data is recommended. This workflow can be run as shell scripts from the command line.

## 5 Discussion

Fetal functional MRI is an emerging field with great potential to improve understanding of human brain development. The number of papers published in this area has seen a 5-fold increase since 2010. Methodologies for processing these complex data sets, however, have not kept pace, and the absence of a standard publicly available processing pipeline for these data has been especially notable. Here we address this gap and present a solution to the most cumbersome processing step, localization and extraction of the fetal brain from surrounding tissue for each volume of a functional time series. This is a necessary step in processing human fetal fMRI data and until now, it has been a rate-limiting factor in accurate, automated processing of fetal fMRI BOLD timeseries data. This model was built by pairing a set of 1,241 hand-drawn fetal 3D brain masks with a deep learning algorithm, a U-Net convolutional neural network. Pipeline code, and documentation are made available through GitHub (https://github.com/saigerutherford/fetal-code). Training, validation, and test data sets (raw volumes, hand drawn masks, and auto-masks) are available through request to Moriah Thomason. It is hoped that release of an easy-to-use, efficient, validated auto-masking pipeline will reduce barriers for new labs to enter this area, while also providing experienced labs opportunities for further optimization of their independently developed approaches.

Recent pioneering work by Salehi and colleagues (2017) established that deep learning approaches can be effective in fetal structural brain segmentation. However, fetal functional imaging presents a distinct set of constraints, and therefore requires a different solution. In particular, the inherently lower resolution and contrast of functional timeseries data, and the 4-dimensional nature of the data (360 3D volumes per subject), make this a more challenging problem. Here we applied CNN methods to the largest fetal fMRI data set reported to date, 207 fetuses, and derived a novel method for accurate and reliable segmentation of the fetal brain from surrounding maternal tissues within a fraction of a second, 94% accuracy in 0.2 seconds. These encouraging results are partially attributable to the large set of manually traced human fetal fMRI masks used to train the CNN, consistent with studies showing the correlation between CNN performance and the size of the training data [48, 49].

The auto-masking model exhibited signs of strong generalizability, which is particularly important given known biases in CNN models trained on imaging datasets from a single site/population [50, 51]. First, the trained CNN correctly classified data at two held out sets with performance, 94% and 89%, very similar to the training data. This finding suggests that the CNN is robust to variations in experimental procedures, scanner settings, and populations studied. In addition, the training images were drawn from a wide fetal age-range, which should also enhance generalizability across fetal samples encompassing different ages.

Our aim in this project goes beyond brain segmentation; we sought to construct an automated version of a previously manual full preprocessing pipeline that is standardized but flexible, and readily deployable across multiple data sources. Thus, our pipeline begins with an auto-masking step, then leverages existing algorithms (FSL, AFNI) that assist with applying the auto-masks, frame to frame alignment, normalization to a user-defined template, and smoothing. The user is referred to publicly available multi-age fetal brain templates [18], and is able to easily configure the tool to modify or eliminate steps. The flexibility of our code allows for potential users to mix and match the portions of this pipeline they wish to use. For example, a user could choose an alternative realignment algorithm as the first step then apply the auto-masking step. The open construction of the tool will allow incorporation of future processing advances, such as surface-based registration and additive motion correction strategies.

Of note, large scale, often multi-center, projects are becoming the new norm, and these require validated, standardized processing pipelines of the kind that we have developed. The Developing Human Connectome Project provides just one example of a large-scale study that includes a fetal functional MRI component [52, 53], and many more large-scale fetal fMRI initiatives will likely emerge in the coming years.

Our work has several limitations. First, deep learning methods perform classification in high dimensional space, and consequently, results can be a “black box” with little opportunity for interpretation of axes [54]. However, this limitation should be viewed in the context of our goal: to automatically perform a task that can take trained individuals many hours to perform manually. As such, we are less interested in understanding the mechanisms of computer-based brain masking, and instead focus on algorithm performance in out of sample data sets.

Another limitation is that we use direct warping, or normalization, of functional data to a group-averaged anatomical template. It is not clear that alternatives would improve registration significantly, but one might expect registration to subject-specific anatomy, then to template space, to be a preferred approach. The challenge of this approach is that obtaining high-quality subject-specific high-resolution anatomical images presents a different set of challenges that have been addressed elsewhere [55]. Prior studies of fetal anatomical development [56] demonstrate that even when trained experts apply the most advanced techniques to these data, there is still significant data loss and image blurring where motion effects, image artifacts, or lack of tissue contrast compromise data quality. This example extends to other parts of the pipeline, where alternative optimized preprocessing strategies could be used. However, the objective for this work is not to serve as a final fetal functional MRI preprocessing endpoint, but as a backbone upon which further development can follow.

A final limitation is that ours is not a fully automated pipeline as it requires human supervision and quality checking at several stages, which in turn requires a certain quantity of time and level of expertise from the human supervisor. Fortunately, however, the level of involvement, and associated expertise required, required is limited, and includes looking for overt errors when viewing processed images as a continuous movie, which takes approximately 1-5 minutes per functional run. Assuming the full time for running this pipeline is 15-30 minutes, including human effort, this is a 60-fold time reduction over prior methods, with manual tracing in particular requiring extensive time and substantial expertise [4, 47].

In sum, in this work, we leverage deep learning methods in the largest sample of fetal fMRI data published to date to address the challenging brain segmentation problem in fetal fMRI. We unite our novel auto-masking tool with other preprocessing steps to initialize the first complete open-source solution to preprocessing raw fetal functional MRI timeseries data.

## 6 Acknowledgements

The authors thank Sahi Karra, Nedda Elewa, Tarek Bazzi, Tahir Khan, Nourhan Hamadi, Bryan Turman, Allison Li, Imran Sheikh, and Sophia Neuenfeldt for their time spent manually generating fetal brain masks, Pavan Jella for assistance in data acquisition, and Lauren Grove for the mentoring and advice while writing this manuscript. The authors also thank participant families who generously shared their time.

## 7 Author Contributions

Conceptualization: SR, MA; Methodology: SR, PS, MA, JH,; Formal Analysis: SR, PS, MA; Data Curation: SR, JH, DS; Writing – Original Draft: SR, PS, JH, MT, CS; Writing – Reviewing and Editing; SR, PS, MA, JH, MV, JW, DS, MT, CS; Visualization: SR, JH; Supervision: MV, JW, MT, CS; Funding Acquisition: MT.

